# Targeting IL-6 reduces IgM levels and tumor growth in Waldenström macroglobulinemia

**DOI:** 10.1101/592642

**Authors:** Weiguo Han, Stephan J. Matissek, David A. Jackson, Brandon Sklavanitis, Sherine F. Elsawa

## Abstract

The tumor microenvironment (TME) plays an important role in cancer and plays a role in resistance to therapy. In Waldenström macroglobulinemia (WM), a B-cell malignancy characterized by the overproduction of a monoclonal IgM protein, the TME plays an important role in WM biology by secreting cytokines that promote malignant phenotype. In previous work, we have shown that TME-IL-6 promotes WM cell growth and IgM secretion in WM. Tocilizumab/Actemra is an anti-IL-6R antibody, which can competitively block IL-6 binding to the IL-6R. We investigated the efficacy of Tocilizumab in a preclinical mouse model of WM that considers the role of the TME in disease biology. Hairless SCID mice were subcutaneously implanted with BCWM.1 or RPCI-WM1 and bone marrow stromal cells. Groups of mice were treated with Tocilizumab or control antibody three times/week for 5 weeks and the effect on tumor burden and disease biology were evaluated. Although Tocilizumab had no effect on mice survival, there was a reduction in tumor growth rate in mice injected with RPCI-WM1 cells treated with Tocilizumab (p=0.0394). In mice injected with BCWM.1 + stromal cells, there was a significant reduction in human IgM secretion in mice sera with Tocilizumab treatment (p=0.0099). There was no significant change in mice weight suggesting Tocilizumab induced no toxicities to the mice. Taken together, our data suggests that administration of Tocilizumab to tumor bearing mice, results in a significant reduction in tumor volume and IgM secretion. Therefore, the evaluation of the role of Tocilizumab in WM patients may provide therapeutic efficacy by reducing IgM production and slowing the rate of tumor growth.

## Introduction

Waldenström macroglobulinemia (WM) is a subtype of non-Hodgkin lymphoma (NHL) characterized by infiltration of the bone marrow with lymphoplasmacytic cells[1]. Despite WM being an indolent lymphoma, WM cells secrete very high levels of a monoclonal immunoglobulin M (IgM) protein, which is associated with symptoms such as anemia, serum hyperviscosity syndrome and peripheral neuropathy[2]. At initial diagnosis, many WM patients do not require immediately clinical intervention unless disease symptoms, mostly associated with hyperviscosity syndrome are evident[2–4]. In the past decade, several significant enhancements were made in our understanding of WM biology. This has led to the evaluation and introduction of several new therapeutic options for WM patients. Current therapies used for the treatment of WM patients mainly focus on targeting cancer cells directly using combination therapies. Rituximab-containing therapies are a standard of care in the United States. This includes therapies such as R-CHOP (Cyclophosphamide, doxorubicin, vincristine and prednisone plus rituximab). Ibrutinib, a BTK inhibitor, is the only FDA approved therapy for WM. However, it is administered indefinitely and can cause adverse reactions such diarrhea, thrombocytopenia, rash, atypical bleeding, among other symptoms[4]. However, despite evidence for a role of the tumor microenvironment (TME) in promoting WM cell growth, survival and IgM secretion[5], there have been little studies investigating targeting the TME as a therapeutic strategy for WM patients.

Despite the progress made to understand disease biology, like most other neoplasms, WM remains an incurable disease and ultimately patients succumb to disease progression. The TME plays an important role in the development and progression of WM and has been shown to play a protective role in resistance to therapy[5–7]. In fact, the cross-talk between malignant cells and cells in the TME favors disease progression and promotes IgM secretion. In previous studies, we and others have shown that IL-6 from the TME promotes IgM secretion and cell growth via binding to the IL-6R on WM cells[8–11]. Therefore, targeting IL-6/IL-6R signaling may provide therapeutic benefit for WM patients, particularly those with high levels of IgM in their serum.

Tocilizumab/Actemra is an anti-IL-6R antibody, which can competitively block IL-6 binding to the IL-6Ra. Tocilizumab has been administered or investigated in several clinical settings in patients with several inflammatory-mediated diseases including Castleman’s disease, Rheumatoid arthritis, Systemic juvenile idiopathic arthritis, Crohn’s disease, Giant cell arteritis, Systemic sclerosis, Systemic lupus erythematosus and multiple sclerosis[12]. These studies indicate that administration of Tocilizumab can be used as a novel therapy to block IL-6 in these chronic inflammatory conditions and suggest its potential role in other diseases in which IL-6 plays a role. Furthermore, several studies have investigated the addition of anti-IL-6 therapy in multiple myeloma (MM) with enhanced results over chemotherapeutic regimens[13–15]. However, despite investigations of anti-IL-6 therapy in autoimmune and malignant disorders, no studies to date have investigated its therapeutic efficacy in WM.

In this study, we report the efficacy of targeting the TME with Tocilizumab in a preclinical mouse model of WM that considers the role of the TME in disease biology. We show that Tocilizumab reduced tumor growth and IgM secretion and suggest it may provide therapeutic benefit to WM patients.

## Results

### Targeting IL-6 in the WM TME does not affect survival

Because bone marrow stromal cells in the WM TME are an important source of IL-6, we subcutaneously implanted hairless SCID mice with BCWM.1 cells or RPCI-WM1 cells and HS-5 stromal cells at a ratio of 5:1 onto the right flank of mice. This allowed us to examine the role of paracrine IL-6 from bone marrow stromal cells, which have been shown to play an important role in malignant B cells in WM[5, 8, 9, 16, 17], on WM cells *in vivo*. Upon tumor appearance, we treated mice with either Tocilizumab or IgG control antibody every other day for 5 weeks (**Figure 1A**) and examined the effect of therapy on mice survival. In accordance with Institutional IACUC, mice were euthanized when tumors reached 2 cm in any dimension or when tumors became ulcerated. We found that Tocilizumab treatment did not affect overall survival in mice xenografted with BCWM.1 or RPCI-WM1 and stromal cells (**Figure 1B**).

**Figure 1.**
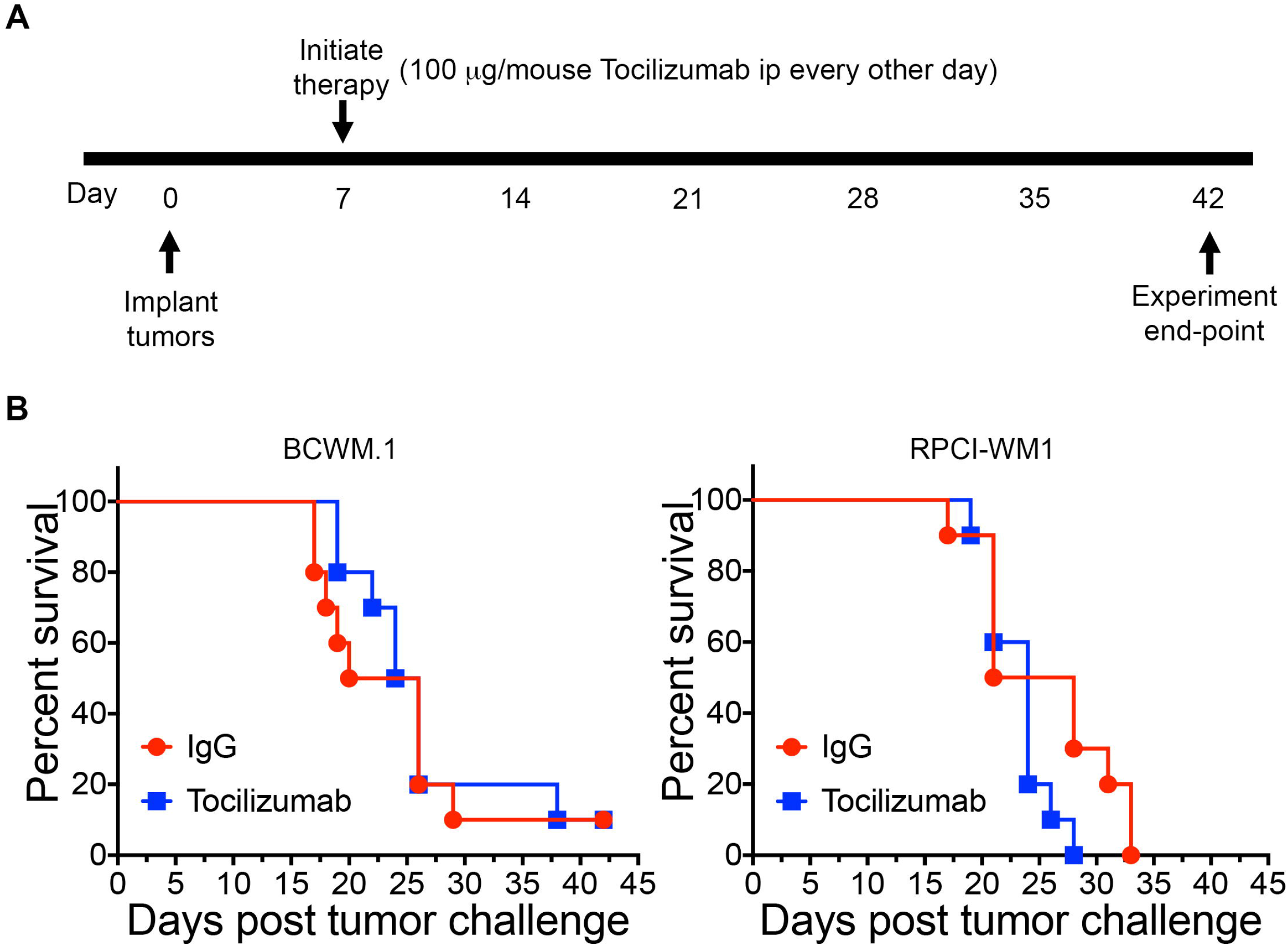
Tocilizumab therapy does not affect overall survival. (A) Hairless SCID mice (n=10 mice/group) were subcutaneously injected with 10 x 10^6^ BCWM.1 or RPCI-WM1 cells + HS-5 stromal cells (5:1 ratio). Upon tumor appearance (day 7), groups of mice were treated with either Tocilizumab or Control antibody (IgG) at 100μg/mouse i.p every other day for a total of 5 weeks. (B) Mice were monitored and survival was reported.

### Targeting IL-6 with Tocilizumab reduces tumor growth and IgM secretion

Tumor growth was monitored and recorded 3 times/week. When we examined the rate of tumor growth (tumor growth relative to the size of the first recorded tumor), we found a significant reduction in tumor growth rate in mice implanted with RPCI-WM1 and stromal cells, and treated with Tocilizumab (p=0.0394) (**Figure 2A**). Although the overall tumor growth rate was not statistically significant in mice implanted with BCWM.1 cells and stromal cells, the rate of tumor growth was slower in Tocilizumab treated mice (**Figure 2A**). Interestingly, when we examined actual tumor volume and tumor growth rate in mice implanted with BCWM.1 and stromal cells at day 14 (when all mice were alive), we found a significant reduction in tumor growth rate (p=0.0306) (**Figure 2B**). We also found a reduction in the actual tumor volume in this group at day 14, although this did not reach statistical significance (p=0.057) (**Figure 2B**). This is consistent with our previous reports on the effect of IL-6 on malignant cell growth in WM, where IL-6 induced a modest increase in WM cell proliferation in this indolent lymphoma[8, 9].

**Figure 2.**
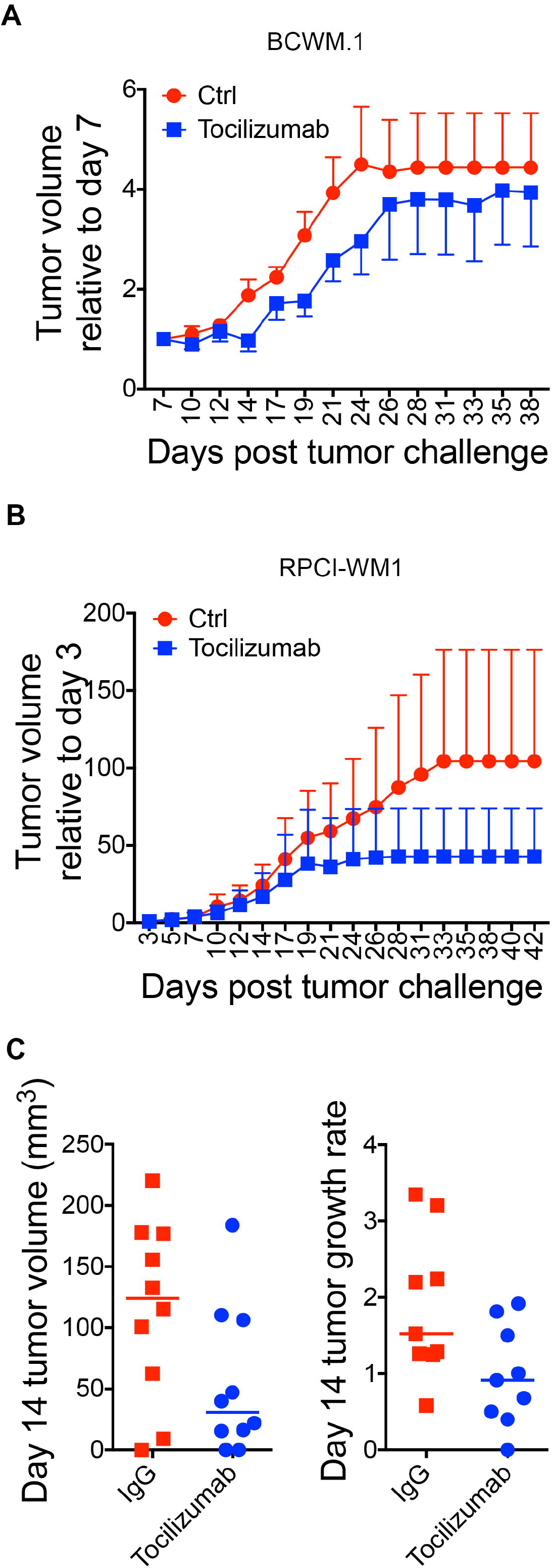
Targeting the TME with Tocilizumab reduces tumor growth. (A) Relative tumor growth rate in BCWM.1 (left; n=10/group) and RPCI-WM1 (p=0.0394)(right; n=10/group) (both with stromal cells) xenografted mice treated with either Tocilizumab or IgG control. The Y-axis indicates tumor volume relative to first recorded tumor size. (B) Tumor volume (left; p=0.057) and relative tumor growth (right, p=0.0306) on day 14 in mice xenografted with BCWM.1 cells and stromal cells.

The role of IL-6 in normal and malignant B cell biology is well established[9, 11, 18–21]. IL-6 has been shown to promote immunoglobulin (Ig) secretion in normal B cells [21] and malignant B cells[9–11, 18–20]. We have previously shown that IL-6 promotes IgM secretion in WM[8, 9]. Therefore, we examined the effect of IL-6 therapy on human IgM secretion in mice sera. Consistent with our previous reports, we found a significant reduction in human IgM secretion in mice sera in groups of mice xenografted with BCWM.1 cells and stromal cells, treated with Tocilizumab (p=0.0029) (**Figure 3**). However, in mice xenografted with RPCI-WM1 cells and stromal cells, there was no reduction in IgM secretion with Tocilizumab treatment (**Figure 3**). The RPCI-WM1 tumors were significantly larger (928.8 +/− 599.6 mm^3^ in control mice) than BCWM.1 xenografted mice (115.1 +/− 73.23 mm^3^ in control mice) prior to euthanasia of mice in either group. Furthermore, of the 3 WM cell lines that are currently available, RPCI-WM1 cells secrete the highest levels of IgM (data not shown).

**Figure 3.**
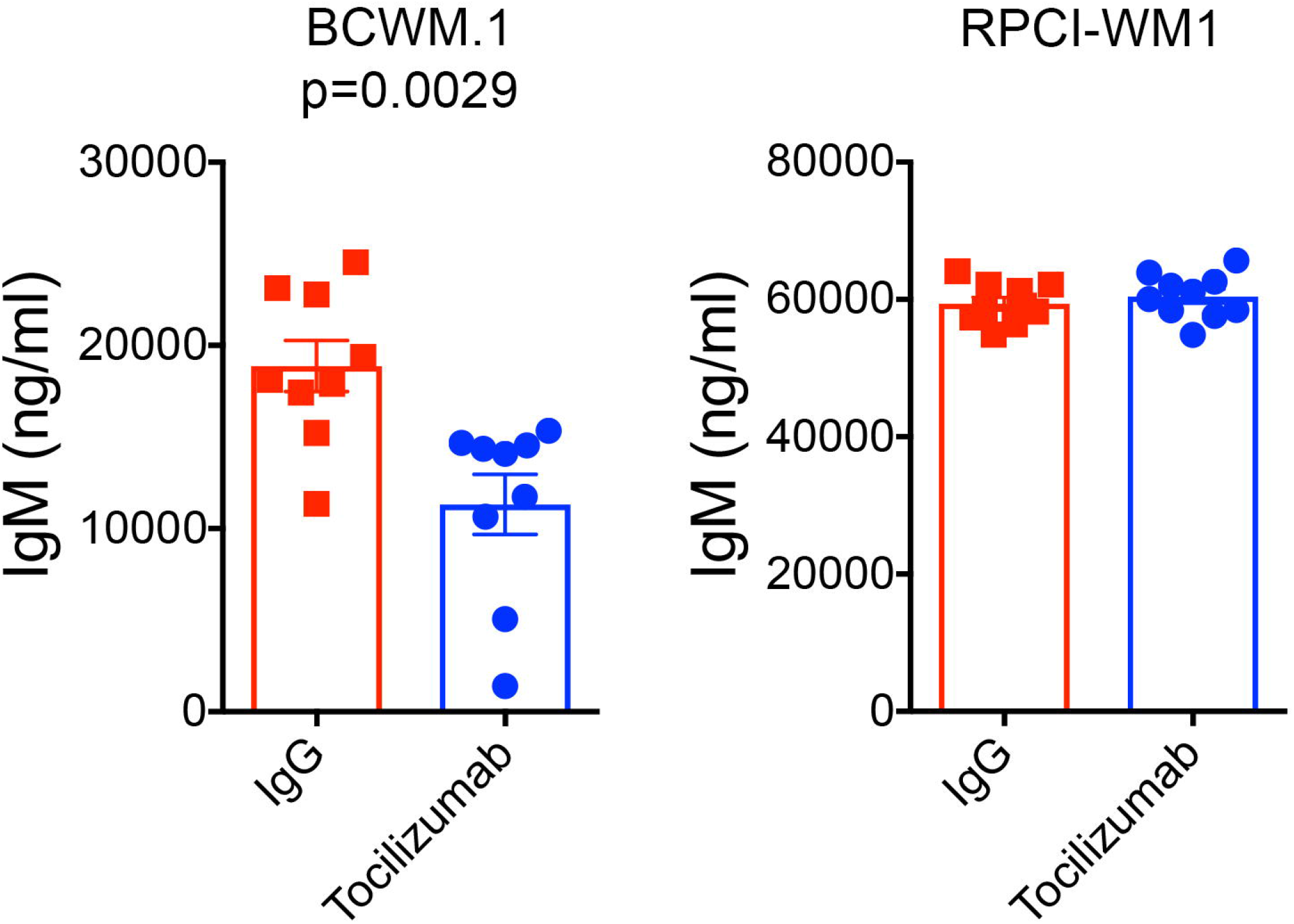
Tocilizumab reduces human IgM secretion in mice xenografted with BCWM.1 cells and stromal cells. Serum was harvested from mice xenografted with BCWM.1 (left; p=0.0029) and RPCI-WM1 (right) cells upon euthanasia and used to quantify human IgM levels in mice sera. IgM levels were determined using a human IgM ELISA.

### Targeting IL-6 with Tocilizumab does not induce toxicity

To examine potential therapy-induced toxicities, we monitored the weights of mice three times/week. Tocilizumab treated mice did not differ from control mice in weight in RPCI-WM1 xenografted mice (**Figure 4A**). In BCWM.1 xenografted mice, there was a significant (p=0.005) difference in mice weights between Tocilizumab treated mice and control mice (**Figure 4A**). However, when we examined individual mice within each group, the data indicates that control mice consistently increased their weight, while the majority (7/10 mice) of Tocilizumab-treated mice either maintained their weight or increased it (**Figure 4B**). Taken together, these results suggest that targeting IL-6 in the TME reduced tumor growth rate and IgM secretion while having no toxic effects on mice.

**Figure 4.**
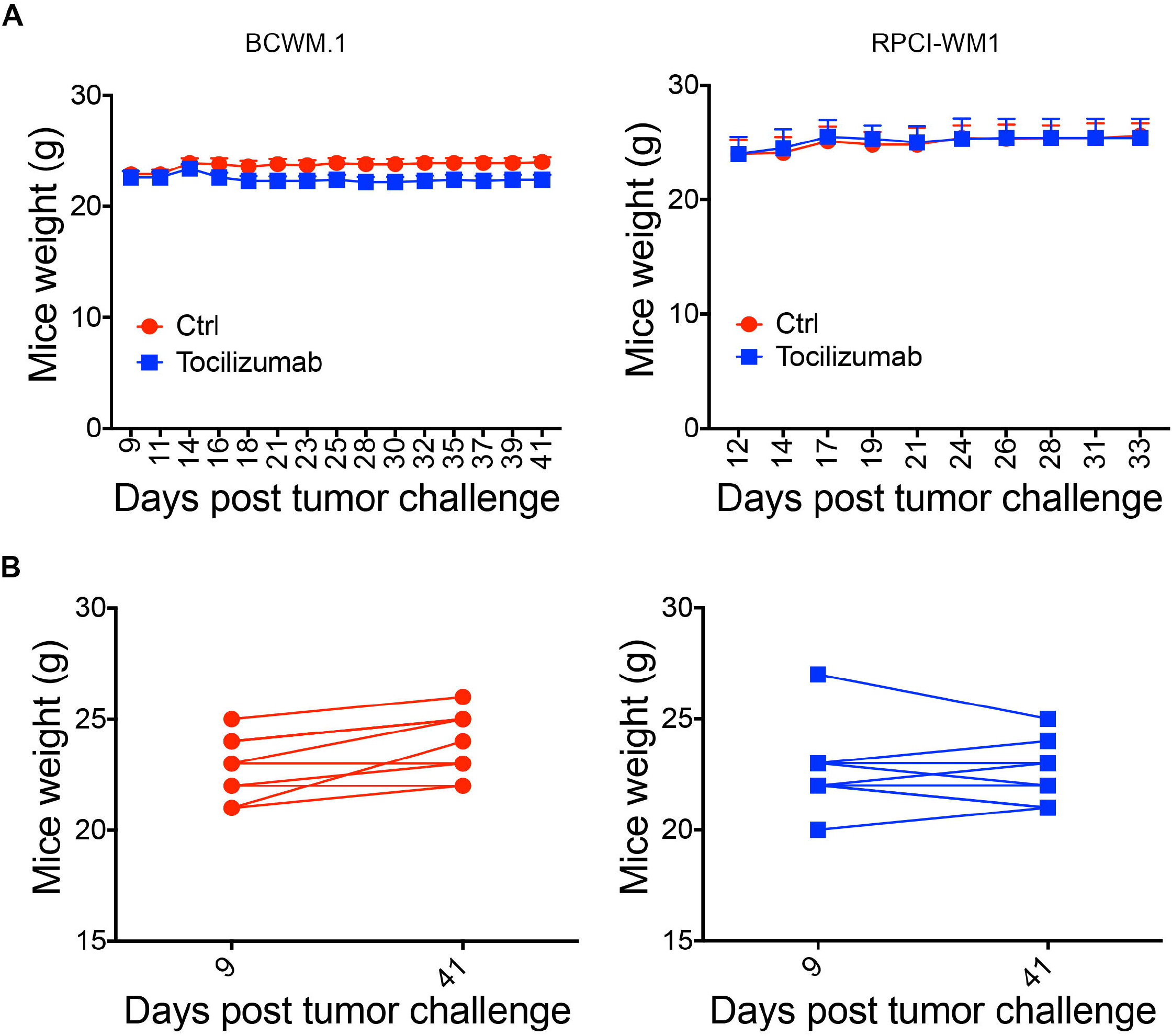
Tocilizumb is not toxic to mice. (A) Mice weights were monitored 3x/week and recoded to determine potential treatment toxicity for BCWM.1 (p=0.005) and RPCI-WM1 xenografted mice. (B) Individual mice weights on day 9 (first recorded weight) and day 42 (experiment end point) for mice xenografted with BCWM.1 cells and stromal cells.

We performed immunohistochemical staining of tumor biopsies with H&E and found a similar cellular morphology composed of small round cells in the 2 mice treatment groups (**Figure 5**). This is consistent with B cell morphology and suggests growth of WM cells to form these tumors.

**Figure 5.**
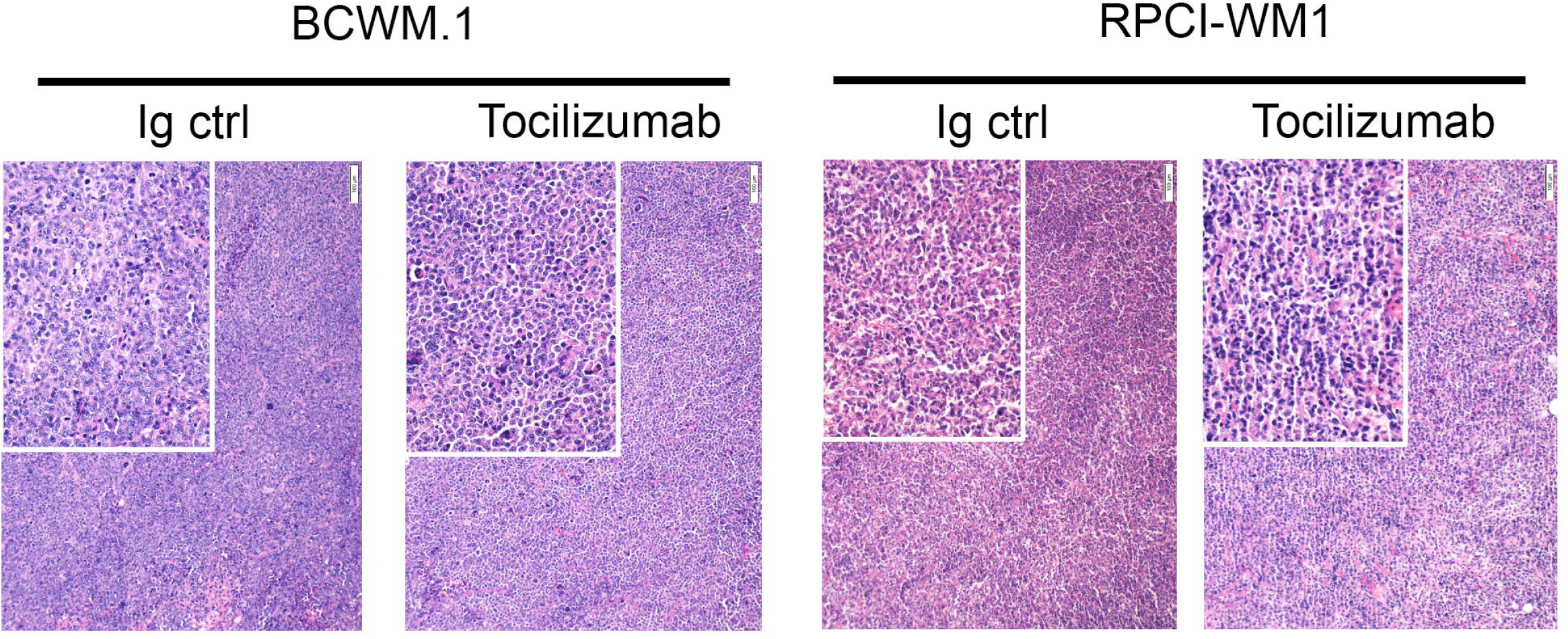
Histology of tumors post euthanasia. H & E staining for tumors harvested from mice treated with Tocilizumab and control (BCWM.1 left, RPCI-WM1 right). Tumor samples were harvested and immediately fixed in 4% formaldehyde. Five μm sections were then stained with H & E as described in the methods.

## Discussion

The role of the bone marrow TME in WM and other B cell malignancies is well documented[6, 16, 22–24]. Interactions between malignant B cells and stromal cells in the TME play an essential role in regulating malignant B cell biology including cell growth, cell survival and Ig secretion[5, 8, 24]. In previous work, we have shown that IL-6 levels are elevated in WM patients and IL-6 from human bone marrow stromal cells can promote WM cells growth and IgM secretion[8, 16]. This finding led us to investigate the efficacy of blocking IL-6 to block the interaction between the TME and WM cells. Our results show that targeting IL-6 in the WM TME did not affect the survival in two WM cell line models investigated (**Figure 1B**). This finding is not surprising, as IL-6 does not induce apoptosis of WM cells [9]. Rather, the role of IL-6 in WM was shown to promote IgM secretion and WM cell growth[8, 9]. Consistent with this role, we did find a reduction in tumor growth rate when tumor bearing mice were treated with Tocilizumab (**Figure 2**). These results indicate that targeting IL-6 in the TME in WM may slow the rate of tumor growth. Since WM remains an incurable disease, a reduction in IgM levels may provide a much needed symptomatic relief for patients. Future studies combining targeting of IL-6 with therapies that induce apoptosis of WM cells, may prove to be effective; with IL-6 therapy slowing tumor growth and another therapy targeting the malignant cells. Currently, Ibrutinib is the only drug approved by the Food and Drug Administration (FDA) for WM patients. However, several other therapies are used as monotherapies or in combination and include Bendamustine, Rituximab, Dexamethasone, Fludarabine, Chlorambucil, Everolimus, among others[1]. An examination of the role of Tocilizumab therapy in combination with these therapies may allow a dose reduction in these therapies and therefore alleviate some of the side effects associated with each therapy.

One of the hallmarks of WM is the overproduction of monoclonal IgM[8–10, 17, 25]. Our results show that tumor-bearing mice treated Tocilizumab had reduced levels of human IgM secretion in mice serum (**Figure 3**). Interestingly, RPCI-WM1 tumor-bearing mice treated with Tocilizumab had similar levels of IgM as control mice. The 2 WM cell lines used in this study are derived from different WM patients[26, 27]. Interestingly, RPCI-WM1 cells secrete the highest levels of IgM among the available WM cell lines (data not shown), raising the possibility that the rate of increase in IgM levels in patient may correlate with the efficacy of Tocilizumab monotherapy. Therefore, an examination of the efficacy of IL-6 therapy in WM patients is necessary to evaluate its effect on IgM levels and tumor growth.

Taken together, these data suggest that administration of Tocilizumab monotherapy to tumor bearing mice results in a reduction in tumor burden and IgM secretion, despite the presence of IL-6 from bone marrow stromal cells in the TME. Therefore, administration of an anti-IL-6 therapy such as Tocilizumab to WM patients may provide therapeutic efficacy by targeting the TME to reduce IgM production and slow[6, 16, 22–24] the rate of tumor growth.

## Materials and methods

### Cells and reagents

The BCWM.1 cells[27, 28] were kindly provided by Dr. Steve Treon (Dana Farber Cancer Institute, Boston, MA), the RPCI-WM1 cells[26, 28] were kindly provided by Dr. Chanan-Khan (Mayo Clinic, Jacksonville, FL) and HS-5 cells were purchased from ATCC (Manassas, VA). WM cells were maintained in RPMI and HS-5 cells in DMEM, all supplemented with 10% FBS and antibiotics/antimycotics as previously published[8–10, 17, 29].

### Mice

Hairless SCID mice (male; 6-8 weeks old; Charles River, Wilmington, MA) were purchased and allowed to acclimate for 2 weeks. Mice were then subcutaneously implanted with BCWM.1 cells (10 x 10^6^) and HS-5 cells (2 x 10^6^) (5:1 ratio) as previously published[8]. Mice were cared for and handled in accordance with institutional and National Institutes of Health guidelines, after obtaining IACUC approval. Mice were treated with 100 μg/mouse either Tocilizumab or control antibody (Genentech, South San Francisco, CA) in a total volume of 100 μl injected via intraperitoneal route every other day for a total of 5 weeks (**Figure 1A**). Tumors were measured (using digital calipers; Fisher Scientific, Waltham, MA) and mice were weighed 3 times/week. Upon euthanasia, tumor samples and sera were harvested and stored for later use.

### IgM ELISA

Human IgM in mice sera was quantified using human IgM ELISA (Bethyl laboratories, Inc., Montgomery, TX) following manufacturer’s recommendations, as previously published[8] using ELISA plates (Nunc Maxisorp, Fisher Scientific). ELISA plates were developed using the Turbo TMB-ELISA (Fisher Scientific) and the reaction was stopped by addition of 2N H2SO4. Results were quantified using a plate reader (Molecular Devices, Palo Alto, CA) and data was analyzed using SoftMax Pro 7.0.2 software.

### Statistical analysis

The last observed outcome data for actual tumor volume, tumor volume relative to baseline and mice weights were compared between treatment groups using a Student’s T-test with the assumption of unequal variance. The mixed effect model with the assumption of random intercepts and slopes for the trajectory lines for each mouse were also fitted to compare the slope of the growth of the outcome measures between treatment groups. The fixed interaction term of the number of days from baseline by treatment indicator were tested for the difference of two arms. All analysis were conducted within tumor type (BCWM.1 or RPCI-WM1).

## Abbreviations

TME: tumor microenvironment
WM: Waldenstrom macroglobulinemia
NHL: Non-Hodgkin Lymphoma
IL-6R: interleukin 6 receptor

## Authorship Contributions

WH, performed the research and wrote the paper; SJM, DAJ and BS performed experiments; SFE oversaw all aspects of the study and wrote the paper.

## Acknowledgements

The authors acknowledge the histology core of the New Hampshire Veterinary Diagnostic Lab (NHVDL) at the University of New Hampshire for tumor sample processing and H & E staining. We also thank Hong Chang from Tufts University for help with statistical analysis.

## References

1. Kapoor P, Paludo J, Vallumsetla N, Greipp PR. Waldenstrom macroglobulinemia: What a hematologist needs to know. Blood Rev. 2015; 29: 301–19. doi: 10.1016/j.blre.2015.03.001.

2. Ansell SM, Kyle RA, Reeder CB, Fonseca R, Mikhael JR, Morice WG, Bergsagel PL, Buadi FK, Colgan JP, Dingli D, Dispenzieri A, Greipp PR, Habermann TM, et al. Diagnosis and management of Waldenstrom macroglobulinemia: Mayo stratification of macroglobulinemia and risk-adapted therapy (mSMART) guidelines. Mayo Clin Proc. 2010; 85: 824–33. doi: 10.4065/mcp.2010.0304.

3. Gertz MA. Selecting Initial Therapy for Newly Diagnosed Waldenstrom Macroglobulinemia. J Clin Oncol. 2018: JCO2018793273. doi: 10.1200/JCO.2018.79.3273.

4. Gertz MA. Waldenstrom macroglobulinemia treatment algorithm 2018. Blood Cancer J. 2018; 8: 40. doi: 10.1038/s41408-018-0076-5.

5. Jalali S, Ansell SM. Bone marrow microenvironment in Waldenstrom’s Macroglobulinemia. Best Pract Res Clin Haematol. 2016; 29: 148–55. doi: 10.1016/j.beha.2016.08.016.

6. Leleu X, Jia X, Runnels J, Ngo HT, Moreau AS, Farag M, Spencer JA, Pitsillides CM, Hatjiharissi E, Roccaro A, O’Sullivan G, McMillin DW, Moreno D, et al. The Akt pathway regulates survival and homing in Waldenstrom Macroglobulinemia. Blood. 2007. doi:

7. Roccaro AM, Sacco A, Husu EN, Pitsillides C, Vesole S, Azab AK, Azab F, Melhem M, Ngo HT, Quang P, Maiso P, Runnels J, Liang MC, et al. Dual targeting of the PI3K/Akt/mTOR pathway as an antitumor strategy in Waldenstrom macroglobulinemia. Blood. 2010; 115: 559–69. doi: 10.1182/blood-2009-07-235747.

8. Elsawa SF, Almada LL, Ziesmer SC, Novak AJ, Witzig TE, Ansell SM, Fernandez-Zapico ME. GLI2 transcription factor mediates cytokine cross-talk in the tumor microenvironment. J Biol Chem. 2011; 286: 21524–34. doi: 10.1074/jbc.M111.234146.

9. Elsawa SF, Novak AJ, Ziesmer SC, Almada LL, Hodge LS, Grote DM, Witzig TE, Fernandez-Zapico ME, Ansell SM. Comprehensive analysis of tumor microenvironment cytokines in Waldenstrom macroglobulinemia identifies CCL5 as a novel modulator of IL-6 activity. Blood. 2011; 118: 5540–9. doi: 10.1182/blood-2011-04-351742.

10. Jackson DA, Smith TD, Amarsaikhan N, Han W, Neil MS, Boi SK, Vrabel AM, Tolosa EJ, Almada LL, Fernandez-Zapico ME, Elsawa SF. Modulation of the IL-6 Receptor alpha Underlies GLI2-Mediated Regulation of Ig Secretion in Waldenstrom Macroglobulinemia Cells. J Immunol. 2015; 195: 2908–16. doi: 10.4049/jimmunol.1402974.

11. Hatzimichael EC, Christou L, Bai M, Kolios G, Kefala L, Bourantas KL. Serum levels of IL-6 and its soluble receptor (sIL-6R) in Waldenstrom’s macroglobulinemia. Eur J Haematol. 2001; 66: 1–6. doi:

12. Tanaka T, Narazaki M, Kishimoto T. Anti-interleukin-6 receptor antibody, tocilizumab, for the treatment of autoimmune diseases. FEBS Lett. 2011; 585: 3699–709. doi: 10.1016/j.febslet.2011.03.023.

13. Hunsucker SA, Magarotto V, Kuhn DJ, Kornblau SM, Wang M, Weber DM, Thomas SK, Shah JJ, Voorhees PM, Xie H, Cornfeld M, Nemeth JA, Orlowski RZ. Blockade of interleukin-6 signalling with siltuximab enhances melphalan cytotoxicity in preclinical models of multiple myeloma. Br J Haematol. 2011; 152: 579–92. doi: 10.1111/j.1365-2141.2010.08533.x.

14. Rossi JF, Fegueux N, Lu ZY, Legouffe E, Exbrayat C, Bozonnat MC, Navarro R, Lopez E, Quittet P, Daures JP, Rouillé V, Kanouni T, Widjenes J, et al. Optimizing the use of anti-interleukin-6 monoclonal antibody with dexamethasone and 140 mg/m2 of melphalan in multiple myeloma: results of a pilot study including biological aspects. Bone Marrow Transplantation. 2005; 36 771. doi: 10.1038/sj.bmt.1705138.

15. Shah JJ, Feng L, Thomas SK, Berkova Z, Weber DM, Wang M, Qazilbash MH, Champlin RE, Mendoza TR, Cleeland C, Orlowski RZ. Siltuximab (CNTO 328) with lenalidomide, bortezomib and dexamethasone in newly-diagnosed, previously untreated multiple myeloma: an open-label phase I trial. Blood Cancer J. 2016; 6: e396. doi: 10.1038/bcj.2016.4.

16. Elsawa SF, Ansell SM. Cytokines in the microenvironment of Waldenstrom’s macroglobulinemia. Clin Lymphoma Myeloma. 2009; 9: 43–5. doi: 10.3816/CLM.2009.n.010.

17. Han W, Jackson DA, Matissek SJ, Misurelli JA, Neil MS, Sklavanitis B, Amarsaikhan N, Elsawa SF. Novel Molecular Mechanism of Regulation of CD40 Ligand by the Transcription Factor GLI2. J Immunol. 2017; 198: 4481–9. doi: 10.4049/jimmunol.1601490.

18. French JD, Tschumper RC, Jelinek DF. Analysis of IL-6-mediated growth control of myeloma cells using a gp130 chimeric receptor approach. Leukemia. 2002; 16: 1189–96. doi:

19. DuVillard L, Guiguet M, Casasnovas RO, Caillot D, Monnier-Zeller V, Bernard A, Guy H, Solary E. Diagnostic value of serum IL-6 level in monoclonal gammopathies. Br J Haematol. 1995; 89: 243–9. doi:

20. Hirano T. Interleukin 6 (IL-6) and its receptor: their role in plasma cell neoplasias. Int J Cell Cloning. 1991; 9: 166–84. doi:

21. Kishimoto T. Interleukin-6: from basic science to medicine--40 years in immunology. Annu Rev Immunol. 2005; 23: 1–21. doi:

22. Burger JA, Peled A. CXCR4 antagonists: targeting the microenvironment in leukemia and other cancers. Leukemia. 2009; 23: 43–52. doi: 10.1038/leu.2008.299.

23. He B, Chadburn A, Jou E, Schattner EJ, Knowles DM, Cerutti A. Lymphoma B cells evade apoptosis through the TNF family members BAFF/BLyS and APRIL. J Immunol. 2004; 172: 3268–79. doi:

24. Burger JA, Ghia P, Rosenwald A, Caligaris-Cappio F. The microenvironment in mature B-cell malignancies: a target for new treatment strategies. Blood. 2009; 114: 3367–75. doi: 10.1182/blood-2009-06-225326.

25. Elsawa SF, Novak AJ, Grote DM, Ziesmer SC, Witzig TE, Kyle RA, Dillon SR, Harder B, Gross JA, Ansell SM. B-lymphocyte stimulator (BLyS) stimulates immunoglobulin production and malignant B-cell growth in Waldenstrom macroglobulinemia. Blood. 2006; 107: 2882–8. doi: 10.1182/blood-2005-09-3552.

26. Chitta KS, Paulus A, Ailawadhi S, Foster BA, Moser MT, Starostik P, Masood A, Sher T, Miller KC, Iancu DM, Conroy J, Nowak NJ, Sait SN, et al. Development and characterization of a novel human Waldenstrom macroglobulinemia cell line: RPCI-WM1, Roswell Park Cancer Institute - Waldenstrom Macroglobulinemia 1. Leuk Lymphoma. 2013; 54: 387–96. doi: 10.3109/10428194.2012.713481.

27. Ditzel Santos D, Ho AW, Tournilhac O, Hatjiharissi E, Leleu X, Xu L, Tassone P, Neri P, Hunter ZR, Chemaly MA, Branagan AR, Manning RJ, Patterson CJ, et al. Establishment of BCWM.1 cell line for Waldenstrom’s macroglobulinemia with productive in vivo engraftment in SCID-hu mice. Exp Hematol. 2007; 35: 1366–75. doi:

28. Drexler HG, Chen S, Macleod RA. Would the real Waldenstrom cell line please stand up? Leuk Lymphoma. 2013; 54: 224–6. doi: 10.3109/10428194.2012.727418.

29. Elsawa SF, Novak AJ, Grote D, Konopleva M, Andreeff M, Witzig TE, Ansell SM. CDDO-imidazolide mediated inhibition of malignant cell growth in Waldenstrom macroglobulinemia. Leuk Res. 2008; 32: 1895–902. doi: 10.1016/j.leukres.2008.03.033.

